# Inhibition of adrenergic β1-AR/Gαs signaling promotes cardiomyocyte proliferation through activation of RhoA-YAP axis

**DOI:** 10.1101/2021.10.20.465083

**Authors:** Masahide Sakabe, Michael Thompson, Nong Chen, Mark Verba, Aishlin Hassan, Richard Lu, Mei Xin

## Abstract

The regeneration potential of the mammalian heart is incredibly limited, as cardiomyocyte proliferation ceases shortly after birth. β-adrenergic receptor (β-AR) blockade has been shown to improve heart functions in response to injury; however, the underlying mechanisms remain poorly understood. Here we inhibited β-AR signaling in the heart using a cardiomyocyte specific β1-adrenergic receptor (β1-AR) blocker (metoprolol) to examine its role in heart maturation and regeneration at the neonatal stage. We found that metoprolol robustly enhanced cardiomyocyte proliferation and promoted cardiac regeneration post myocardial infarction, resulting in reduced scar formation and improved cardiac function. Moreover, the increased cardiomyocyte proliferation was also induced by the genetic deletion of *Gnas*, the gene encoding G protein alpha subunit (Gαs), a downstream effector of β-AR. Genome wide transcriptome analysis revealed that the cardiomyocytes of β-blocker treated and *Gnas* cKO hearts maintained an immature proliferating status even at the young-adult age, and that the loss of Gαs function enhanced the activity of the Hippo-effector YAP, which is associated with immature cardiomyocyte proliferation. Moreover, the increased YAP activity is modulated by RhoA signaling. Our pharmacological and genetic studies reveal a previously unrecognized β1-AR-Gαs-YAP signaling axis for regulating cardiac regeneration. These results suggest that inhibiting β-AR-Gαs signaling promotes the regenerative capacity and extends the cardiac regenerative window in mice by activating YAP-mediated transcriptional programs. Thus, targeting β-AR-Gαs signaling may serve as a novel therapeutic target for the treatment of ischemic heart.

## Introduction

The capability of regeneration and repair in response to cardiac injury in the adult mammalian heart is limited. Neonatal mouse hearts retain regenerative potential following cardiac injury up to 7 days after birth^1-3^. Changes after birth such as metabolic state, oxygen level, cardiomyocyte structure and maturity, hormones, and polyploidy, are among the factors contributing to the loss of the regenerative potential in the heart^4-10^. For instance, the postnatal metabolic shift from glycolysis to fatty acid oxidation or aerobic respiration-mediated oxidative DNA damage can lead to cardiomyocyte cell cycle arrest postnatally ^6, 7, 10^. In addition, signaling pathways such as Hippo, Neuregulin, ERBB2, Agarin, and thyroid hormone have been shown to regulate cardiac regeneration^4, 11-14^.

The evolutionarily conserved Hippo signaling pathway is known as a pivotal regulator of organ size and cell proliferation^15, 16^. It consists of a series of kinases, including MERLIN, MST1/2, and LATS1/2, that phosphorylate the downstream effectors YAP and TAZ, preventing their nuclear translocation and activation of target gene expression^17, 18^. Activation of YAP in the embryonic heart, either through the loss of *MST1/2, Sav1*, or forced expression of a constitutively active form of YAP, induces cardiomyocyte proliferation and increases heart size^19, 20^. Moreover, activation of YAP in adult hearts improves cardiac function and reduces scar formation after myocardial infarction (MI)^21-23^, suggesting that YAP activation promotes cardiomyocyte regeneration even in the adult mouse heart. However, it remains unknown what upstream signaling cues regulate the Hippo signaling pathway for cardiac regeneration.

The Hippo pathway can be activated by several molecular signals, including G protein-coupled receptors (GPCRs), cell-cell interaction, and alterations in cytoskeletal dynamics^24-26^. Beta-adrenergic receptors (β-ARs), members of GPCRs that couple to a stimulatory G protein alpha-subunit (Gαs), are essential components of the sympathetic nervous system^27^. Stimulation of the β-ARs activates Gαs activity leading to an increase in intracellular cAMP levels and the subsequent activation of protein kinase A (PKA), which can result in increased heart rate and contractility^28, 29^. Overexpression of Gαs-protein leads to increased myocardial collagen content and fibrosis with variable hypertrophy in mice^30, 31^, suggesting an important role of Gαs in controlling cardiac contractility or hypertrophic response in the heart. Furthermore, activation of Gαs by epinephrine inactivates YAP and inhibits cell growth in various cell lines^24^. This presents an opportunity to manipulate cardiomyocyte proliferation through inhibition of the β-AR signaling pathway.

Inhibition of β-AR by β-adrenergic receptor blockade (β-blockers) has been shown to improve the survival and symptoms in heart failure patients^32, 33^. Gene variants in *GNAS*, which encodes the Gαs protein, are associated with β-blocker-related survival or risk in patients after coronary artery bypass grafting^34^. A recent study suggested that a non-selective β-blocker (propranolol) increased the number of cardiomyocytes in neonatal mice^35^. Since propranolol inhibits both the β1-AR (expressed in the heart) and β2-AR (present in vascular and non-vascular smooth muscle cells in non-heart tissues), it is not suitable for patients with diabetes or bronchospasm. The 2^nd^ generation β1-blocker, metoprolol, selectively binds to the β1-AR receptor on cardiomyocytes^36^; however, its effect on cardiomyocyte proliferation and heart regeneration have not been explored. Furthermore, the mechanisms by which β-AR-Gαs signaling modulates cardiomyocyte proliferation and heart regeneration remains to be defined.

In this study, we show that treatment with the β1-blocker metoprolol, promotes cardiomyocyte proliferation, reduces scar formation, and improves cardiac function after myocardial injury. Inhibition of the β1-AR downstream effector Gαs activity by genetic deletion of *Gnas* enhances cardiomyocyte proliferation by activating YAP activity through its nuclear localization. Thus, our study demonstrates that β-AR-Gαs signaling represses the regenerative capacity of neonatal cardiomyocytes by inhibiting YAP-activated transcriptional programs. Inhibition of β-AR-Gαs signaling extended the cardiac regenerative window, suggesting a potential therapeutic role of the β1-blocker metoprolol in cardiac regeneration after heart injury.

## Materials and Methods

### Mouse experiments

All animal experiments were performed with the approval of the Institutional Animal Care and Use Committee of Cincinnati Children’s Hospital Medical Center. Mouse lines harboring the *Gnas* and *Yap* floxed alleles have been described previously^19, 37^. The α-Myosin heavy chain (*Myh6*)-Cre and *Myh6-MerCreMer* mice were obtained from Jeff Robbins and Jeffery D. Molkentin, respectively^38, 39^. Tamoxifen (Sigma) was dissolved in 90% sunflower oil (Sigma)/10% ethanol and stored at -20°C. Prior to intraperitoneal (IP) injection (50 mg/kg per day), the tamoxifen solution was heated at 55 °C for 10 min. For 5-ethynyl-2-deoxyuridine (EdU) studies, mice were administered an IP injection of EdU (5 μg/kg) once a day. The β-blocker (metoprolol) was dissolved in saline and injected by IP (2 mg/kg) once a day. C57BL/6J mice were used for the β-blocker treatment studies.

### Myocardial infarction

To induce myocardial infarction (MI) in neonatal mice, we permanently ligated the left anterior descending artery on P7 as previously described^40, 41^. Briefly, mice were anaesthetized with isoflurane and the heart was exposed via thoracotomy through the fourth or fifth intercostal space. An 8-0 nylon suture was tied around the left anterior descending coronary artery (LAD). Subsequently, the chest and skin were closed in layers using 6-0 nylon sutures. The mouse was allowed to recover from surgery on a heating pad. Sham-operated mice underwent the same procedure involving anesthesia and thoracotomy without LAD ligation.

### Echocardiography

Assessment of cardiac function on conscious, non-sedated mice was performed with the Vivo 2100 micro-ultrasound system (VisualSonics). Cardiac function and heart rate were measured on M-mode and doppler images.

### Neonatal rat cardiomyocyte isolation and culture

Neonatal rat cardiomyocyte culture was performed using the neonatal cardiomyocyte isolation kit (Cellutron). P2 neonatal cardiomyocytes were plated on tissue culture dishes pre-coated with SureCoat (Cellutron) at a density of 2×10^5^/cm^2^. After 24 hours, cardiomyocytes were treated with epinephrine (100 μM, Sigma), C3 (1mg/ml, Cytoskeleton), and GO4 (100 μM, provided by Dr. Yi Zheng).

### Mouse cardiomyocyte isolation

The mouse cardiomyocyte isolation was performed as previously described^42^. In brief, P14 hearts were harvested and immediately fixed with 4% PFA at 4 °C overnight. Subsequently, samples were incubated with collagenase B (1.8 mg/ml, Roche) and D (2.4 mg/ml, Roche) for 12h at 37 °C. The hearts were minced to smaller pieces and the procedure was repeated until no more cardiomyocytes were dissociated from the tissue. The digested cardiomyocytes were stained with 4’,6-diamidino-2-phenylindole (DAPI) for nucleation counts. A hemocytometer was used for counting cardiomyocytes.

### Assessment of cardiomyocyte size

The cross-sectional area of cardiomyocytes was assessed using wheat germ agglutinin (WGA) staining. Cryosections were rinsed in PBS and then incubated with WGA conjugated with Alexa Fluor 488 (1:100, Invitrogen). Slides were imaged by Eclipse Ti confocal microscopy with a C2 laser-scanning head (Nikon). ImageJ software (National Institutes of Health) was used to quantify the size of each cell. The area of the digested cardiomyocytes was quantified using ImageJ software based on phase contrast images.

### Histological analysis

Hearts were fixed in 4% paraformaldehyde (PFA) at 4 °C overnight, embedded in paraffin, and sectioned at 5μm thickness. Hematoxylin and eosin (H&E) staining was performed following standard protocol. Masson’s trichrome and Picrosirius red staining was performed according to standard procedures at CCHMC’s pathology core. Fibrotic area was quantified using ImageJ software.

### Immunofluorescence experiments

For immunostaining, hearts were fixed in 4% PFA at 4 °C overnight, embedded in OCT compound (StatLab), and sectioned at 8μm thickness. For PCM1 staining, we used fresh-frozen (non-fixed) samples and sections were fixed in 10% formalin. Sections were blocked with 1% bovine serum albumin (BSA), incubated with primary antibodies against PH3 (rabbit polyclonal, 1:200; Millipore), PCM1 (1:1000; Sigma), cardiac α-actinin (1:200; Sigma), cardiac Troponin T (1:200; Thermo), YAP (1:100; Cell Signaling), Runx1 (1:100; Cell Signaling), smooth muscle α-actin conjugated with AlexaFluor594 (1:200; Sigma), and were further incubated with Alexa Fluor-conjugated secondary antibodies against mouse, rabbit IgG and with DAPI. For EdU staining, neonates were administered an intraperitoneal (IP) injection of 5-ethynyl-2-deoxyuridine (EdU, 5μg/g of mouse body weight). EdU incorporation was assessed using Click-IT EdU system (Invitrogen). Fluorescence images were captured using Eclipse Ti confocal microscopy with a C2 laser-scanning head (Nikon).

### RNA sequencing

RNA from hearts were extracted using Trizol (Invitrogen) followed by purification using RNeasy Mini kit (Qiagen). RNA-seq was performed using two individual animals for control and *Gnas* cKO hearts, or control and β-blocker treated hearts. RNA sequencing was performed by the Center for Medical Genomics, Indiana University School of Medicine. Gene Ontology analysis of gene expression changes was performed using Enrichr^43^ and Gene Set Enrichment Analysis (GSEA) software.

### Real time qPCR

Total RNA was isolated using Trizol according to the manufacture’s protocol. cDNA was synthesized from 500 ng of total RNA using PrimeScript RT Master Mix (Takara). Quantitative real-time PCR (qPCR) was performed using SYBR-Green Master Mix (KAPA) on a StepOnePlus Real-Time PCR system (Applied Biosystems). Values for specific genes were normalized to 18s ribosomal RNA.

### Active RhoA assay

RhoA activity was examined by an effector domain, GST-fusion pull-down protocol, as previously described^44^. Cultured cardiomyocytes or heart tissues were lysed in a lysis buffer containing 1% Triton X-100 and incubated with the glutathione bead-bound GST-Rhotekin. The bead-immobilized GTP-bound RhoA and total RhoA in the lysates were probed by immunoblotting with anti-RhoA antibody (Cell Signaling).

### Western blot analysis

Cultured cardiomyocytes or heart tissues were lysed with 2x sample buffer (BioRad) containing 2-mercaptoethanol and heated for 5 minutes at 95°C. Equal amounts of protein were run on SDS-polyacrylamide gel and transferred to Immobilon-P membrane (Millipore). Membranes were incubated with anti-YAP (Novus), anti-phospho-YAP (Cell Signaling), and anti-GAPDH (Cell Signaling) antibodies at 4°C overnight. Anti-rabbit horse-radish peroxidase (GE healthcare) was used as the secondary antibody, followed by detection with Super Signal West Pico chemiluminescent substrate (Thermo).

### Statistical analysis

All datasets were taken from n ≥ 3 biological replicates. Used animal numbers or group numbers are described in the respective figure legends. Animals were genotyped before the experiments and were caged together and treated in the same way. The experiments were not randomized. We calculated p values with unpaired Student’s t test or analysis of variance (ANOVA) followed by Tukey-Kramer test with Excel (Microsoft Office). P value < 0.05 was considered to represent a statistically significant difference. Data are presented as mean ± SD.

## Results

### β1-blocker treatment promotes neonatal cardiomyocyte proliferation

β1-AR is the main receptor in the heart, and β2-AR is highly expressed in the vascular and non-vascular smooth muscle cells. To investigate whether blockade β1-AR specifically has a significant effect on cardiomyocyte proliferation and cardiac regeneration, we injected the β1-AR blocker metoprolol (hereinafter, referred to as “β-blocker”) in neonatal mice for two weeks starting at postnatal day (P) 1 via daily intraperitoneal injection (IP) (Figure 1a). The heart rate of β-blocker treated mice was significantly reduced and thickening of the myocardial wall of β-blocker treated hearts is evident at P14 (Figure 1b and c). β-blocker treatment induced a significant increase in the heart weight-to-body weight ratio at P14, but not at P7 (Figure 1d and Supplemental Figure 1a). We detected increased proliferating cardiomyocytes, mononucleated cardiomyocytes (a proliferative and regenerative subpopulation of the postnatal heart) and total number of cardiomyocytes in the β-blocker-treated hearts (Figure 1e, 1f and Supplemental Figure 1b-d), while the cardiomyocyte cell size was comparable between control and β-blocker-treated hearts (Figure 1f and Supplemental Figure 1e-f). Moreover, β-blocker treatment promoted cardiomyocyte proliferation at later time points from P14 to P28 (Figure 1g), when the majority of cardiomyocytes are mature, suggesting that β1-AR-selective blocker treatment reactivates cell proliferation of mature cardiomyocytes *in vivo*.

**Figure 1.**
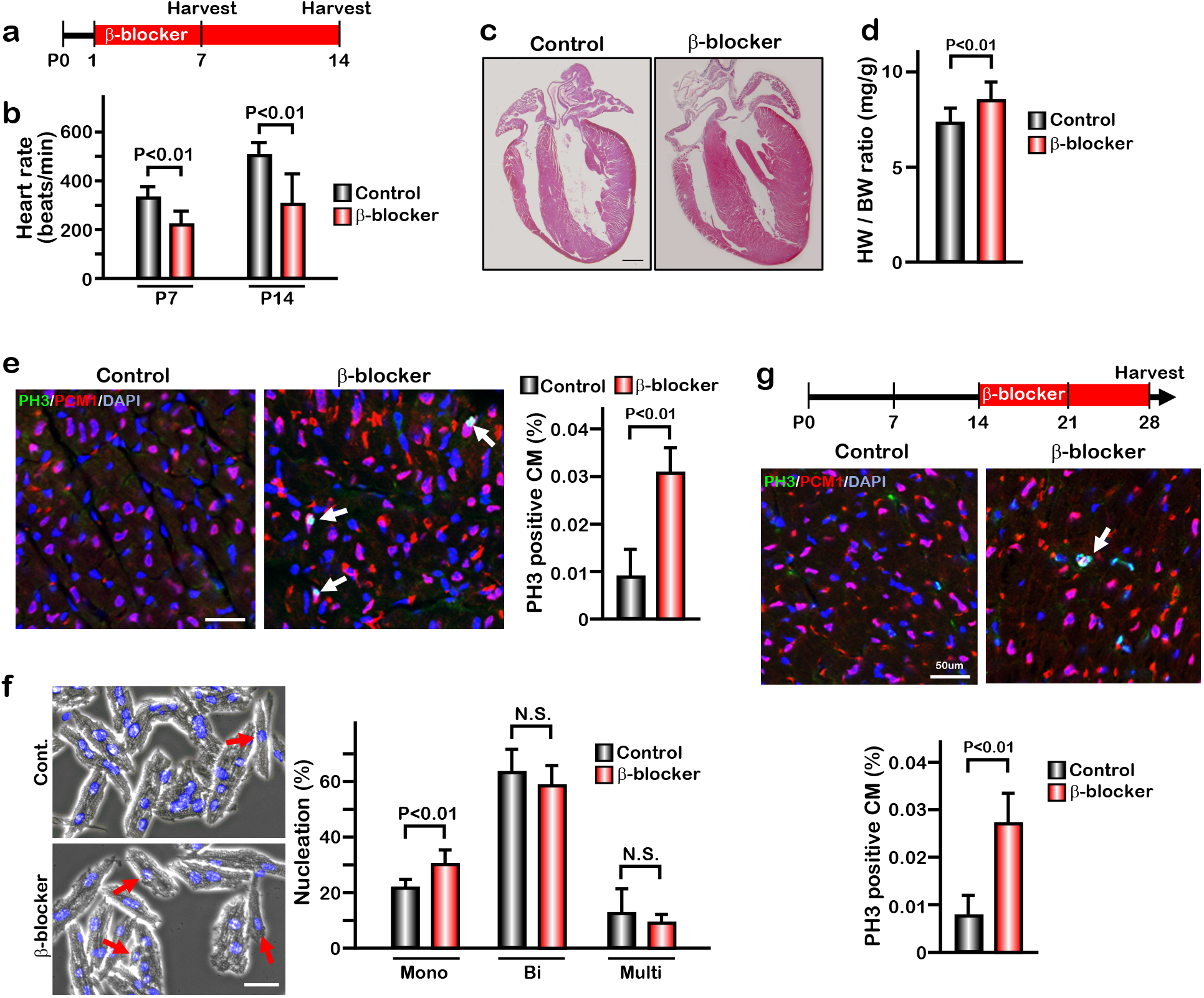
β-blocker (Metoprolol) treatment promotes neonatal cardiomyocyte proliferation. **(a)** Schematic of experimental timeline. **(b)** Heart rate of the saline (control) and β-blocker-treated mice at P7 and P14. Data are as mean ± SD, (P7, n=6; P14, n=10). **(c)** H&E staining of control and β-blocker-treated hearts at P14. Scale bar: 500μm. **(d)** Heart weight (HW) to body weight (BW) ratio at P14. Data are as mean ± SD, n=10. **(e)** Co-immunostaining of PH3 and PCM1 of heart sections from control and β-blocker-treated mice at P14 (left panel). Quantification of PH3 positive cardiomyocytes (right panel). Data are as mean ± SD, n=4. Scale bar: 50μm. **(f)** DAPI staining of isolated cardiomyocytes from P14 control and β-blocker treated hearts. Quantification of Mono-nucleated, Bi-nucleated and multi-nucleated cardiomyocytes in P14 control and β-blocker treated hearts. Data are as mean ± SD, n=4. N.S., not significant. Scale bar: 100μm. **(g)** PH3 and PCM1 co-immunostaining of P28 heart sections of control and mice with daily β-blocker treatment since P14 (upper panel). Quantification of PH3 positive cardiomyocytes (lower panel). Data are as mean ± SD, n=4. Scale bar: 50μm.

### β1-blocker treatment promotes cardiac regeneration after myocardial infarction

To determine whether β-blocker treatment has any effect on neonatal heart regeneration, we induced myocardial infarction by permanent ligation of the left anterior descending coronary artery (LAD) at P7, and treated the mice with β-blocker from P8 until P28 (Figure 2a). Whereas the vehicle treated mice showed loss of heart tissue, extensive scarring and dilation post MI, β-blocker treated mice exhibited significantly reduced left ventricle fibrosis and increased myocardial tissue (Figure 2b and Supplemental Figure 2). Cell-proliferation of cardiomyocytes was upregulated by β-blocker treatment in both the infarct border zone and the remote zone, evidenced by an increase in the number of PH3 positive cardiomyocytes (Figure 2c-e), suggesting that β-blocker treatment promotes neonatal cardiac regeneration by enhancing cardiomyocyte proliferation in the injured hearts. Moreover, echocardiography analysis indicated that β-blocker treatment led to an improvement in cardiac function post MI (Figure 2f-g). Both ejection fraction (EF) and fractional shortening (FS) were decreased in all mice relative to sham control mice 1 week after MI. By 3 weeks post MI, cardiac function was dramatically enhanced in the β-blocker treated hearts (Figure 2f-g). These data indicate that β1-selective blocker treatment in neonatal mice can promote heart regeneration and sustain cardiac functions post MI injury.

**Figure 2.**
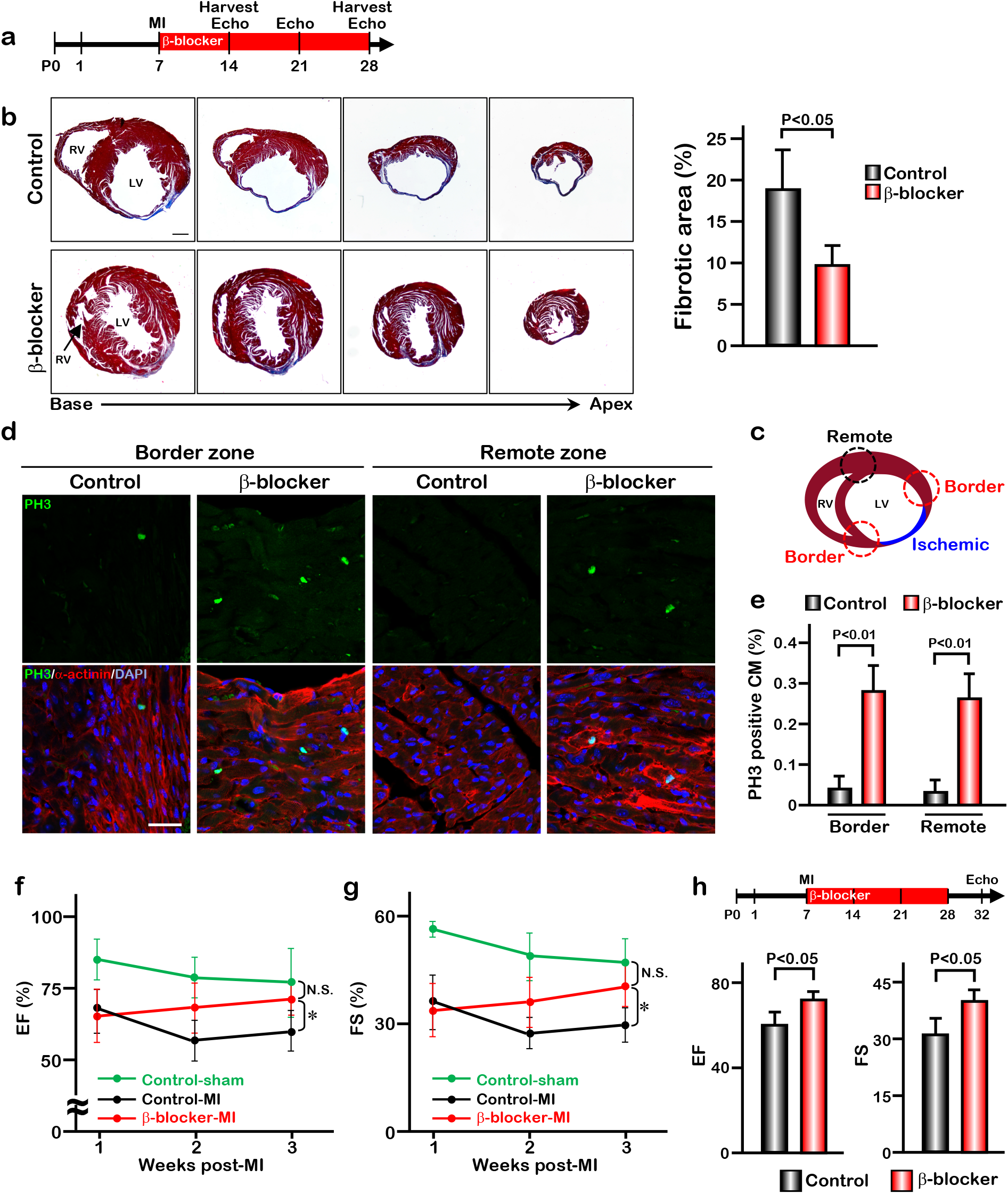
β-blocker treatment promotes cardiac regeneration and cardiomyocyte proliferation following injury in young-adult hearts. **(a)** Schematic of experimental timeline. **(b)** Masson’s trichrome staining of heart sections from control and β-blocker-treated mice 3 weeks post-MI. Scale bar: 500μm. Quantification of the fibrotic areas (Right panel). Data are as mean ± SD, n=4. **(c-e)** PH3 and cardiac α-actinin staining of injured hearts (border and remote zone) treated with saline or β-blocker. Data are as mean ± SD, n=3. Scale bar: 25μm. **(f-g)** Echocardiographic analysis of control and β-blocker-treated mice at 3 weeks post MI surgery. Serial echocardiographic measurements of EF and FS of injured hearts treated with saline (Control) or β-blocker (n=9). *, P<0.01; N.S., not significant. **(h)** P7 mice were subjected to LAD ligation and treated with β-blocker from P8 to P28. 4 days after final β-blocker treatment, the heart function was assessed by echocardiography. Data are as mean ± SD, n=3.

The improved cardiac function in the β-blocker treated mice could be due to a larger volume of blood flow into the ventricle, leading to an increase in the force of contraction associated with a slower heart rate. To rule out this possibility, we assessed cardiac function at 4 days after the last β-blocker treatment. Since metoprolol has a short half-life of 3-7 hours, it should be metabolized during these 4 days. Although the heart rates were comparable between β-blocker treated and saline treated control mice, we found that cardiac function was still significantly improved in the β-blocker treated heart (Figure 2h). Therefore, these results suggest that the β-blocker treatment improves cardiac function by promoting cardiac regeneration in the injured heart.

### Deletion of *Gnas* promotes cardiomyocyte proliferation

β1-AR is associated with the stimulatory G protein (Gαs) and the activated Gα subunit then regulates the down-stream effector molecules such as PKA and cAMP. We hypothesize that blockade of β1-AR promotes cardiomyocyte proliferation through the inhibition of Gαs activity. To test this hypothesis, we deleted *Gnas*, the gene encoding Gαs, in the heart by crossing *Gnas*^*flox/flox*^ mice with *Myh6-Cre* mouse line. The *Gnas*^*flox/flox*^; *Myh6-Cre* (*Gnas* cKO) hearts did not show any abnormal phenotype compared with littermate controls at birth. However, from P7 onwards, *Gnas* cKO hearts were markedly enlarged and the heart weight-to-body weight ratio was significantly increased (Figure 3a-b, Supplemental Figure 3a). Furthermore, enzyme-linked immunosorbent assay (ELISA) showed that cAMP levels were significantly reduced in *Gnas* cKO hearts (Figure 3c), suggesting that Gαs function was reduced in *Gnas* cKO hearts. Consistent with the downregulation of cAMP level, heart rate was also decreased in *Gnas* cKO (Figure 3d).

**Figure 3.**
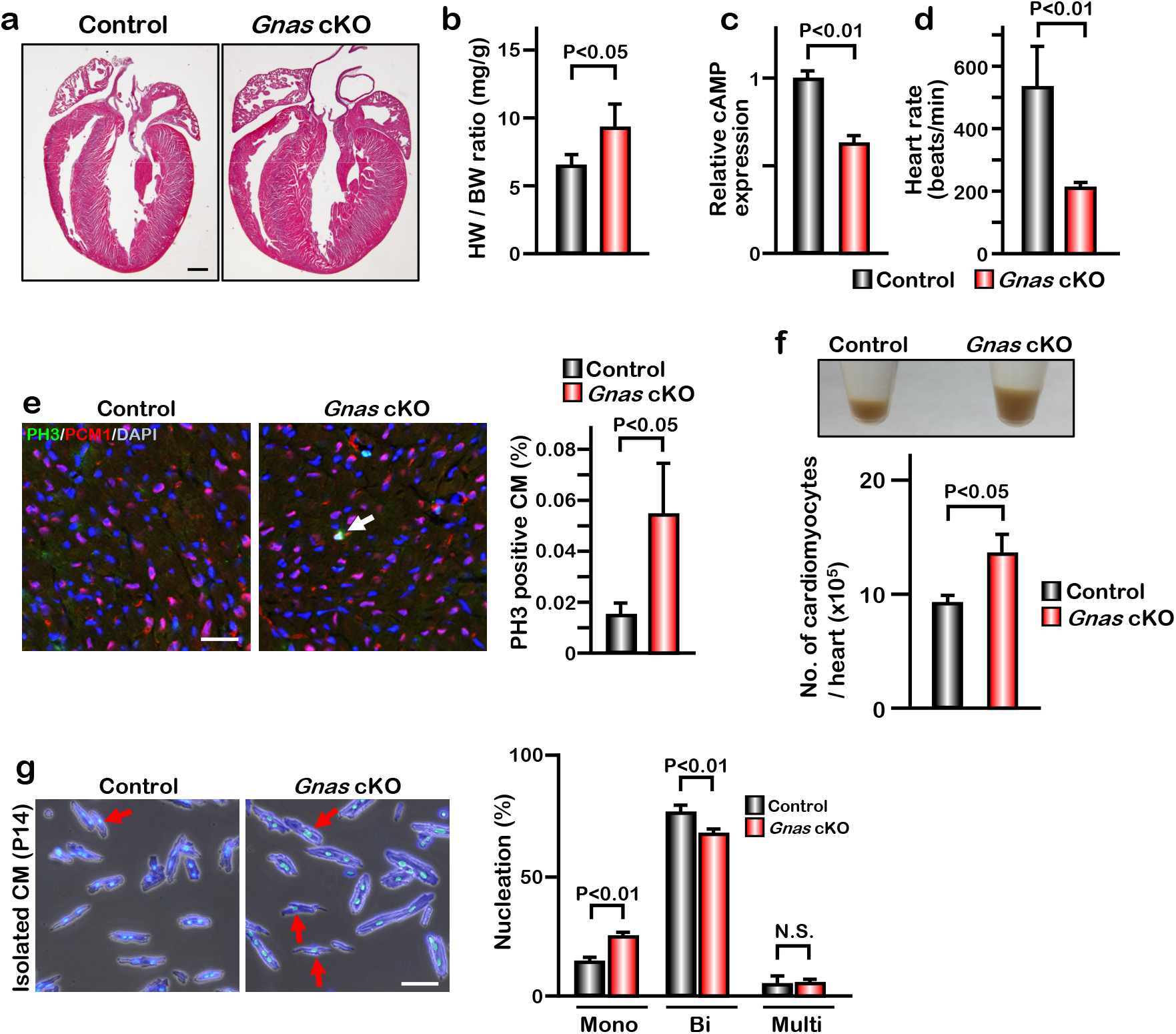
Deletion of *Gnas* promotes cardiomyocyte proliferation. **(a)** Hematoxylin and eosin staining of P7 control and *Gnas* cKO heart sections. Scale bar: 500μm. **(b)** Heart weight (HW) to body weight (BW) ratio of P7 *Gnas* cKO mice. Data are as mean ± SD, n=5. **(c)** Relative cAMP level in P7 control and *Gnas* cKO hearts. Data are as mean ± SD, n=3. **(d)** Heart rate at P14 in control and *Gnas* cKO mice. Data are as mean ± SD, n=5. **(e)** PH3 and PCM1 staining of left ventricle sections of control and Gnas cKO mice at P14 (left panel). Quantification of PH3 positive cardiomyocytes (right panel). Data are as mean ± SD, n=3. Scale bar: 50μm. **(f)** Pellet of cardiomyocytes isolated from a WT and Gnas cKO heart at P14 (upper panel) and quantification of number of isolated cardiomyocytes (lower panel). Data are as mean ± SD, n=3. **(g)** DAPI staining of isolated cardiomyocytes from P14 control and Gnas cKO hearts. Quantification of mono-nucleated, bi-nucleated and multi-nucleated cardiomyocytes in P14 Gnas cKO hearts (right panel). Data are as mean ± SD, n=3. Scale bar: 50μm.

Immunobiological analysis revealed that the deletion of *Gnas* in the heart results in enhanced proliferation of cardiomyocytes evidenced by increased number of a proliferation marker PH3 in PCM1 positive cardiomyocytes (Figure 3e). Further, we dissociated P14 hearts by collagenase and found that the number of cardiomyocytes was significantly increased in *Gnas* cKO hearts (Figure 3f). No significant difference in cardiomyocyte cell size was detected between the control and *Gnas* cKO hearts (Supplemental Figure 3b-c), suggesting that the increased heart size was not caused by cardiac hypertrophy. The percentage of mononucleated cardiomyocytes in *Gnas* cKO hearts was indeed higher than that in control hearts (Figure 3g). These data suggest that *Gnas* ablation leads to increased cardiomyocyte proliferation and heart size but not cardiac cell hypertrophy.

### Inhibition of β1-AR-Gαs signaling promotes metabolic switch from fatty acid oxidation to glycolysis in cardiomyocytes

To determine the potential mechanisms by which inhibition of β-AR-Gαs promotes cardiomyocyte proliferation and cardiac regeneration, we performed RNA-sequencing analysis (RNA-Seq) using RNA isolated from P7 control, *Gnas* cKO, and β-blocker treated hearts. We identified approximately 2000 differentially regulated genes (fold change ≥ 1.2) between control vs. *Gnas* cKO and control vs. β-blocker treated hearts. We found that 1076 and 733 genes were down-regulated in *Gnas* cKO and β-blocker treated hearts, respectively, and that 109 genes overlapped between *Gnas* cKO and β-blocker treated hearts. Gene ontology (GO) analysis using Enrichr^43^ indicated that the expression of genes related to fatty acid metabolism, a major source of energy for mature cardiomyocytes, was down-regulated in *Gnas* cKO and β-blocker treated hearts (Figure 4a). Gene set enrichment analysis (GSEA) also showed that fatty acid metabolism-related genes were down-regulated in *Gnas* cKO and β-blocker-treated hearts (Figure 4b, Supplemental Figure 4a-b). In contrast, the expression of genes related to glycolysis, the metabolic pathway utilized by immature cardiomyocytes and associated with cardiomyocyte proliferation, was upregulated in *Gnas* cKO and β-blocker treated hearts (Supplemental Figure 4a-d). Quantitative PCR (q-PCR) analysis of fatty acid metabolism-related genes confirmed the RNA-seq results (Figure 4c). Moreover, ultrastructural analysis of the heart by electron microscopy (EM) revealed elongated mitochondria in control hearts, whereas mitochondria of *Gnas* cKO hearts were small and round, which is a distinctive phenotype of immature mitochondria, suggesting down-regulation of total energy metabolism in *Gnas* cKO and β-blocker treated hearts. (Figure 4d). Consistent with EM images, the copy number of mitochondrial DNA was less in *Gnas* cKO hearts (Figure 4e), These data suggest that cardiomyocytes in the *Gnas* cKO and β-blocker treated hearts exhibit a characteristic feature of immature cardiomyocytes.

**Figure 4.**
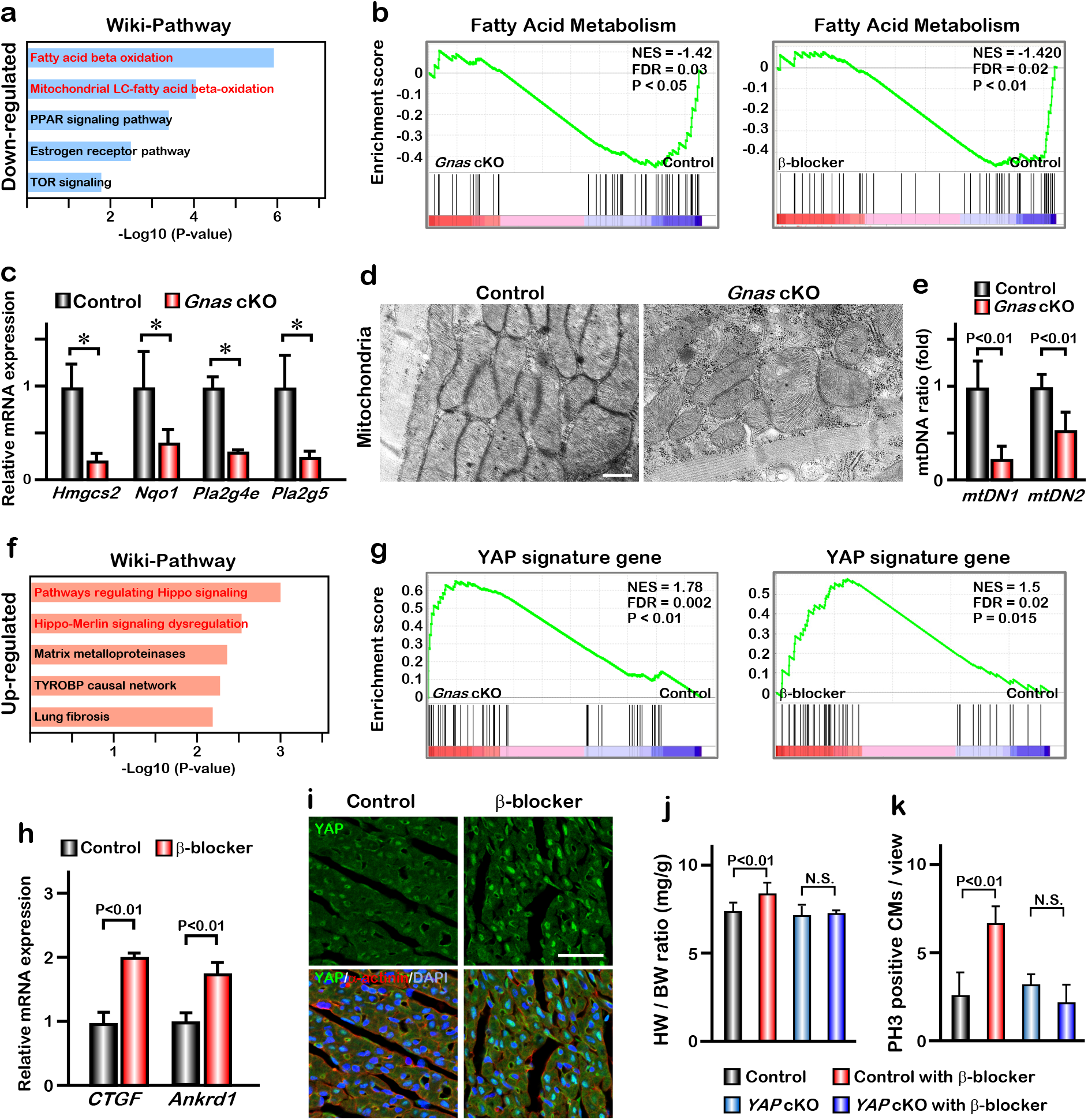
Inhibition of βAR-Gαs signaling leads to elevation of YAP activity in the cardiomyocytes. **(a)** Functional enrichment of GO terms for the common down-regulated genes in *Gnas* cKO and β-blocker treated hearts at P7 (fold change ≤ 0.8). **(b)** GSEA plot shows that fatty acid metabolic genes are down-regulated in *Gnas* cKO and β-blocker treated hearts. **(c)** Q-PCR analysis of fatty acid metabolism related genes in control and *Gnas* cKO hearts at P14. Data are as mean ± SD, n=3. **(d)** Transmission electron microscopy images of mitochondria in ventricular cardiomyocytes of P14 control and *Gnas* cKO hearts. Scale bar: 600nm. **(e)** Q-PCR analysis of mitochondrial DNA in control and *Gnas* cKO hearts at P14. Mitochondrial DNA copy number was normalized to nuclear DNA copy number (mtDN1 vs. H19, and mtDN2 vs. Mx1). Data are as mean ± SD, n=3. **(f)** Functional enrichment of GO term for the up-regulated genes in *Gnas* cKO and β-blocker treated hearts at P7 (fold change ≥ 1.2). **(g)** GSEA plot showed that YAP signature genes are up-regulated in *Gnas* cKO and β-blocker treated hearts. **(h)** Q-PCR analysis of YAP target gene expression, CTGF and Ankrd1, in control and β-blocker treated hearts at P14. Data are as mean ± SD, n=3. **(i)** YAP and cardiac α-actinin immunostaining of heart sections from control and β-blocker treated mice at P7 (n=3). Scale bar: 50μm. **(j)** Heart weight (HW) to body weight (BW) ratio of P14 control (n=15), control with β-blocker (n=13), Yap cKO (n=4), and Yap cKO with β-blocker (n=5) treated mice. Data are as mean ± SD, n=5. N.S., not significant. **(k)** Quantification of the number of PH3 positive cardiomyocytes per view. Data are as mean ± SD, n=4. N.S., not significant.

### Inhibition of β1-AR-Gαs signaling promotes YAP transcriptional activity leading to proliferative immature cardiomyocytes

GO and GSEA analysis indicated that the expression of genes related to the Hippo signaling pathway was increased in *Gnas* cKO and β-blocker treated hearts (Figure 4f-g, Supplemental Figure 4a-b). q-PCR analysis confirmed the upregulation of YAP target genes *CTGF* and *Ankrd1* in *Gnas* cKO and β-blocker treated hearts (Figure 4h). Consistent with the YAP target gene expression, blockade of β1-AR-Gαs signaling by either β-blocker treatment or deletion of *Gnas* induced YAP nuclear translocation (Figure 4i, Supplemental Figure 5a). Furthermore, YAP also translocated to the nucleus when the *Gnas* gene was deleted later at P14 upon tamoxifen treatment in the *Gnas*-*MCM* cKO mouse line (Supplemental Figure 5b). This suggests that Gαs inhibits YAP nuclear localization not only in neonatal but also in postnatal cardiomyocytes. Conversely, when Gαs was activated by epinephrine, an agonist for β-AR, YAP translocated from the nucleus to the cytoplasm at P4 in the control mice; however, this cytoplasmic translocation of YAP did not occur in the *Gnas* cKO hearts (Supplemental Figure 5c), suggesting that β-AR signaling blocks YAP nuclear localization through Gαs activation in the neonatal hearts.

**Figure 5.**
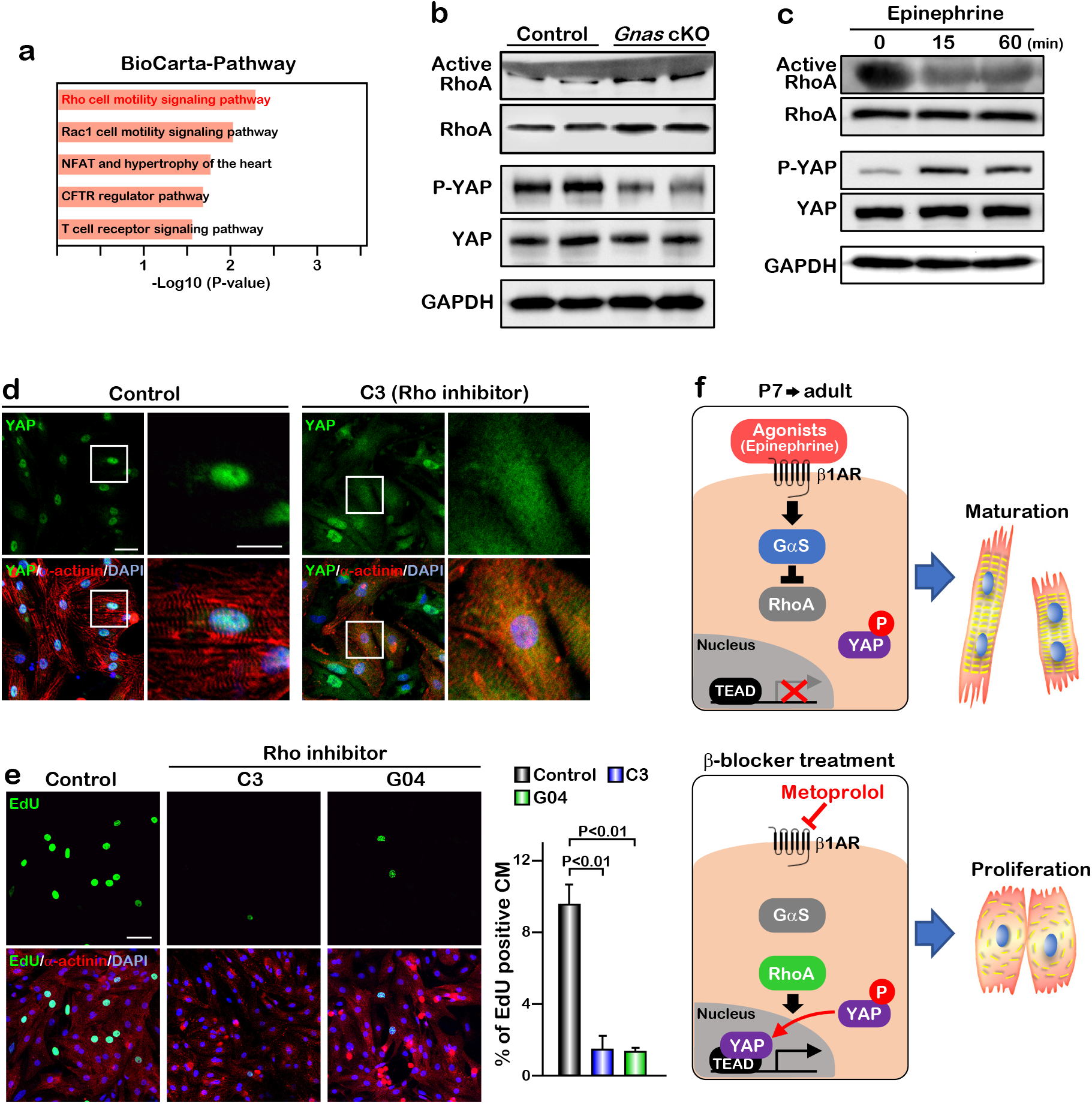
GαS regulates cardiomyocyte proliferation through RhoA mediated YAP activation. **(a)** Functional enrichment of GO terms for the common up-regulated genes. **(b)** Active-RhoA pull-down assay and western blot analysis of P7 control and *Gnas* cKO hearts. **(c)** Active-RhoA pull-down assay and western blot analysis of cultured cardiomyocytes with epinephrine treatment. **(d)** Immunostaining of YAP in cultured cardiomyocytes treated with C3 (Rho inhibitor). Scale bar: 50μm (left panel) and 25μm (right panel). **(e)** EdU incorporation assay on rat neonatal cardiomyocytes treated with Rho inhibitors (left panel). Quantification of EdU-labelled proliferating cardiomyocytes stained with cardiac α-actinin. Data are as mean ± SD, n=3 (right panel). Scale bar: 50μm. **(f)** Model of β1-AR-Gαs signaling regulation of cardiomyocyte proliferation.

### Ablation of YAP abolished the β-blocker induced cardiomyocyte proliferation

We next investigated whether the increased proliferation phenotype seen with β-blocker treatment was caused by an activation of YAP. We performed β-blocker treatment in the cardiac specific *Yap* knockout mice *Yap* ^*flox/flox*^*;aMHCCre* (*Yap* cKO), which did not show any morphological phenotype in hearts at P14, and found that β-blocker treatment did not increase cardiomyocyte proliferation in the *Yap* cKO hearts (Figure 4j-k), suggesting that β-blocker-induced cardiomyocyte proliferation is dependent on YAP functions. Together, our data suggest that Gαs mediates adrenergic signaling to inhibit cardiomyocyte proliferation via inhibition of YAP activity.

### Gαs inhibits cardiomyocyte proliferation through inactivation of the Rho signaling pathway

To gain insight into the molecular mechanisms of how Gαs regulates YAP activity in neonatal cardiomyocytes, we performed pathway analysis based on transcriptomic profiles and found that Rho signaling pathway was activated in *Gnas* cKO and β-blocker treated hearts (Figure 5a). To examine whether Gαs regulates RhoA activity *in vivo*, we performed active-RhoA pull down assay using P7 control and *Gnas* cKO hearts. As expected, RhoA activity was increased while the inactive form of phospho-YAP was dramatically decreased in the *Gnas* cKO heart, compared with littermate controls at P7 (Figure 5b).

To confirm this inhibitory effect of Gαs against RhoA, we stimulated Gαs with epinephrine in cultured neonatal rat cardiomyocytes. Epinephrine treatment resulted in a decrease in RhoA activity and an increase in YAP phosphorylation (Figure 5c). Moreover, when cardiomyocytes were treated with C3 toxin, a Rho inhibitor, a proportion of YAP was translocated from the nucleus to the cytoplasm, while YAP remained in the nucleus in the saline-treated cardiomyocytes (Figure 5d). Treatment with the Rho inhibitors C3 and G04 greatly reduced cardiomyocyte proliferation, as demonstrated by EdU incorporation (Figure 5e). Thus, our data suggest that β1-AR-Gαs signaling negatively regulates cardiomyocyte proliferation through the inhibition of RhoA mediated YAP activity. Given that pharmacological inhibition of Gαs promotes cardiomyocyte proliferation through activation of YAP, β-blocker could be used as a potential therapeutic drug to promote cardiomyocyte proliferation and heart regeneration (Figure 5f).

## Discussion

Since the adult mammalian heart has limited potential for re-entry into the cell cycle, cardiomyocyte replenishment after the loss of cardiomyocytes due to myocardial infarction is insufficient to restore the heart function^45-47^. To repair or improve heart function of the injured heart, several strategies using cellular therapies, such as direct cardiac reprogramming, and noncellular therapies have been published^48-51^. In this study, we identify the β1-adrenergic/Gαs-protein signaling as an inhibitory pathway that restricted the capacity of cardiomyocytes to return to immature proliferative states. We show that pharmacological and genetic inhibition of β1-adrenergic/Gαs-protein reactivate cardiomyocyte proliferation programs and heart regeneration after the regenerative window by activating Hippo-YAP signaling, pointing to a potential therapeutic strategy for inducing cardiac repair in injured heart.

Although neonatal cardiomyocytes have the capacity to proliferate, this regenerative potential is lost in mice one week after birth. During this narrow window, cardiomyocytes undergo one round of DNA synthesis and nuclear division without cytokinesis, which leads to binucleated cardiomyocytes and cell cycle arrest^52^. At the same time, metabolism in cardiomyocytes switches from glycolysis to fatty acid metabolism, and contractile proteins change from embryonic to neonatal isoforms^53-55^. It has been suggested that cardiomyocyte maturation is conversely correlated with its proliferative ability with the increase of DNA content and well aligned contractile structures. Recent studies also demonstrated that oxidative stress after birth is one driver for cell cycle arrest in neonatal cardiomyocytes^6, 7^. Despite these findings, the molecular mechanisms of postnatal cardiomyocyte cell cycle withdrawal are not fully understood. In the present study, we found that Gαs signaling activity was correlated with the loss of proliferative or regenerative ability of neonatal cardiomyocytes, and inhibition of Gαs activity promotes mono-nucleated cardiomyocyte division and cardiac regeneration after the regenerative window. We demonstrated that inhibition of Gαs induces the de-differentiation of mature cardiomyocytes to an immature state and reactivates YAP transcriptional activity to promote cardiac regeneration after myocardial infarction.

It has been shown that cardiac Gαs regulates heart rate and myocardial contractility ^29, 56^. However, the role of Gαs in cardiomyocytes proliferation is still unclear. Our cardiomyocyte-specific *Gnas* KO mice showed an enlarged heart phenotype with an increase in cardiomyocyte proliferation at neonatal stage. Conversely, transgenic mice overexpressing Gαs displayed myocardial hypertrophy with increased myocardial collagen content and fibrosis^30^. Together, these results suggest that Gαs plays an important role in maintaining cardiomyocyte homeostasis.

Several signals such as mechanical and oxidative stress have been identified as regulators of the Hippo signaling pathway^57-60^. YAP is known as a mechanical sensor due to its ability to alter its localization in response to various environmental stimuli, including alternation of cytoskeletal dynamics and blood flow^57, 58, 61, 62^. We found that treatment of cultured cardiomyocytes with a Rho inhibitor hinders YAP activity. This is consistent with a previous study showing that the Rho-mediated pathway promotes nuclear localization of YAP in human embryonic stem cells^24^. At present, it is still unclear whether Rho directly regulates YAP activity in cardiomyocytes. A previous report using a cardiomyocyte-specific RhoA transgene revealed heart rate depression and decreased cardiomyocyte contraction^63^. We found that RhoA activity was gradually downregulated between P0 and P14 (Supplemental Figure 6), and heart rate gradually increased from ∼250 beats/min at P0 to ∼550 beats/min at P14^64^ (Supplemental Figure 7). During this period, YAP translocates from the nucleus to the cytoplasm, suggesting that RhoA may regulate YAP activity through inhibiting cardiomyocyte contraction rate.

β-AR signaling has been shown to increase cardiac output by enhancing heart rate and contractility via activation of Gαs protein^65, 66^. β1-AR is expressed in all the cardiomyocytes with equal distribution between the left and right ventricles. On the other hand, β2-AR is not only expressed in the cardiomyocytes, but also in vascular and non-vascular smooth muscle cells in multiple tissues. Both β1-and β2-AR are coupled to GαS, while β2-AR can be coupled to Gαi/o as well^56^. Stimulation of β1-AR results in adenylyl cyclase-mediated cAMP generation and upregulation of PKA, MAP kinase, and calmodulin-dependent protein kinase II (CAMKII)^66-68^. Clinical studies indicated that β-blockers, especially the β1-AR-specific blocker, metoprolol, improve cardiac function and reduce mortality in patients with heart failure and MI^33^. However, mechanisms underlying the therapeutic effects of β-blockers in heart failure patients are poorly understood. A recent study indicated that in heart failure patients, YAP activity is inactivated by phosphorylation, and that blocking the inhibitory kinase LATS1/2 can reverse heart failure post MI in mice^23^. Similarly, we demonstrate that β-blocker treatment promotes nuclear YAP translocation and cardiac regeneration after MI. Given that YAP activity enhances cardiac regeneration and promotes survival post-MI in our mouse models, activation of Hippo-YAP signaling might be one of the reasons why β-blocker treatment is able to improve heart function in patients.

## Acknowledgements

The authors thank Dr. Jeff Molkentin for insightful discussions and suggestions; Dr. Masayuki Fujii for helpful discussions; Zhifei Xu, Bin Liu, and Hui Sun for technical support; Lingli Xu for Data analysis; Dr. Yi Zheng for the Rho inhibitor GO4; Dr. Eric N. Olson for *Yap* floxed mice; Dr. Lee Weinstein for *Gnas* floxed mice. This work was supported by the National Institutes of Health (Grant HL-132211).

---

**Supplemental Figure 1.**
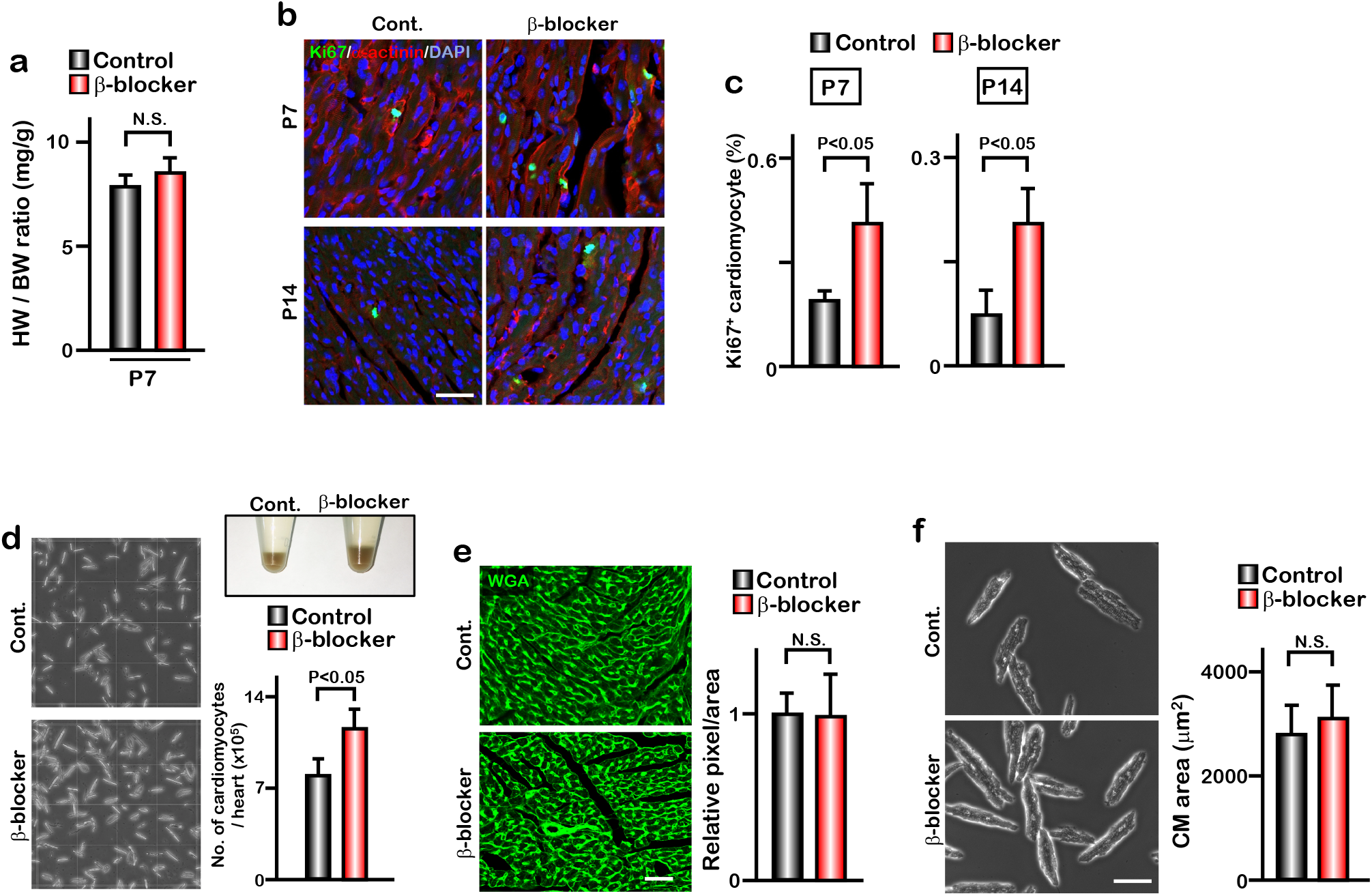
β-blocker treatment promotes cardiomyocyte proliferation after the cardiac regeneration window. **(a)** Heart weight (HW) to body weight (BW) ratio of β-blocker treated heart at P7 (n=10). **(b)** Ki67 and cardiac α-actinin staining of heart sections from control and β-blocker treated mice at P7 and P14. Scale bar: 50μm. **(c)** Quantification of Ki67 positive cardiomyocytes at P7 and P14. Data are as mean ± SD, n=4. **(d)** Images of isolated cardiomyocytes and quantification of number of isolated cardiomyocytes from control and β-blocker treated hearts at P14. Data are as mean ± SD, n=3. Pellets of isolated cardiomyocytes from a whole heart are shown in 1.5 ml tubes. **(e)** Wheat germ agglutinin (WGA)-stained sections of the LV compact zone in hearts of P14 control and β-blocker treated mice (n=3). Scale bar: 50μm. Quantification of relative pixel per area of WGA-positive cardiomyocytes (n=8) (right panel). N.S., not significant. **(f)** Images (left) and quantification (right) of isolated cardiomyocytes from control and β-blocker treated mice. Twenty cardiomyocytes from 3 individual hearts were measured. Scale bar: 100μm.

**Supplemental Figure 2.**
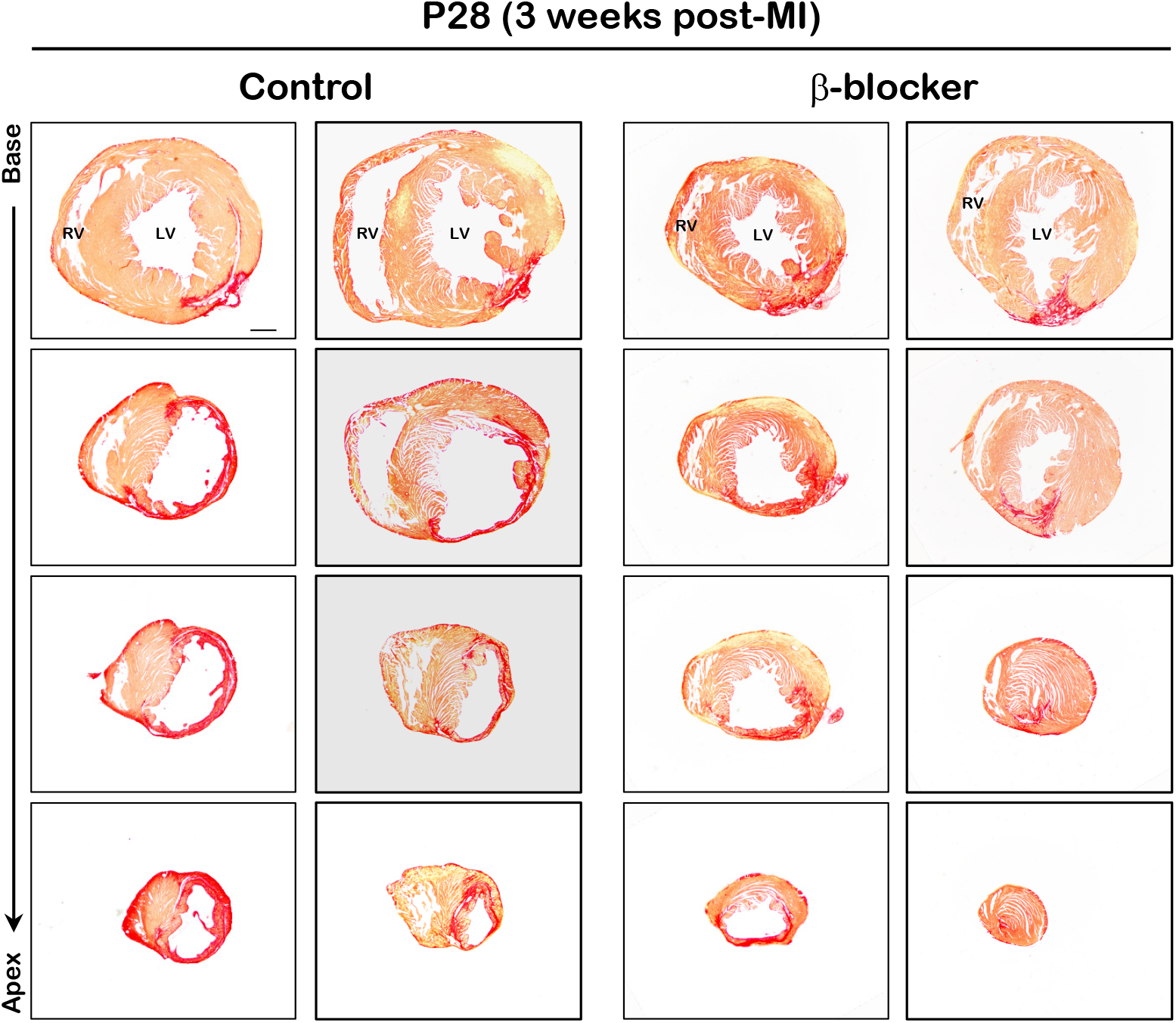
Scar area is reduced in β-blocker treated hearts 3 weeks post MI. Hearts were subjected to LAD ligation at P7 and treated with β-blocker from P8 for 3 weeks. Scar area was analyzed by sirius red staining of transverse sections at P28. Serial sections were cut at 200μm intervals from the site of the ligature to the apex. Red region indicates fibrotic scar area. Scale bar: 500μm.

**Supplemental Figure 3.**
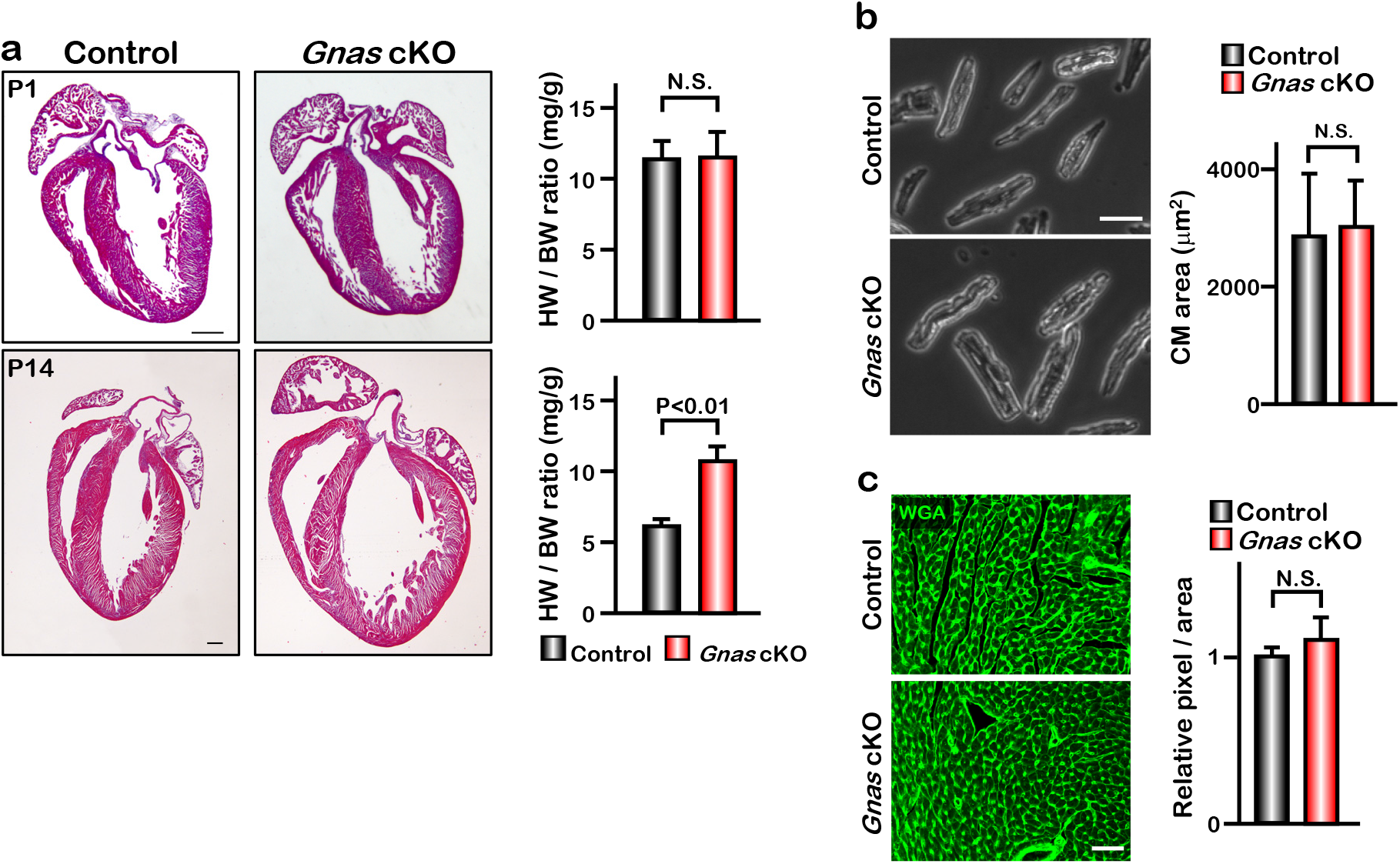
*Gnas* cKO hearts exhibit enlarged phenotype but do not show cardiac hypertrophy. **(a)** H&E staining of control and *Gnas* cKO hearts at P1 (upper panels) and P14 (lower panels) and quantification of heart weight-to-body weight ratio of control and *Gnas* cKO neonates at P1 and P14, respectively. Data are as mean ± SD, n=7. Scale bar: 500μm. **(b)** Isolated cardiomyocytes from control and *Gnas* cKO mice at P14. Scale bar: 100μm. Quantification of the area of isolated cardiomyocytes measured with Image J software (Right panel). Data are as mean ± SD, 20 cardiomyocytes from 3 individual hearts were measured. **(c)** Wheat germ agglutinin (WGA) staining of P14 control and *Gnas* cKO heart sections. Quantification of relative pixel per area of WGA-positive cardiomyocytes (Right panel). N.S., not significant. Data are as mean ± SD, n=8. Scale bar: 50μm.

**Supplemental Figure 4.**
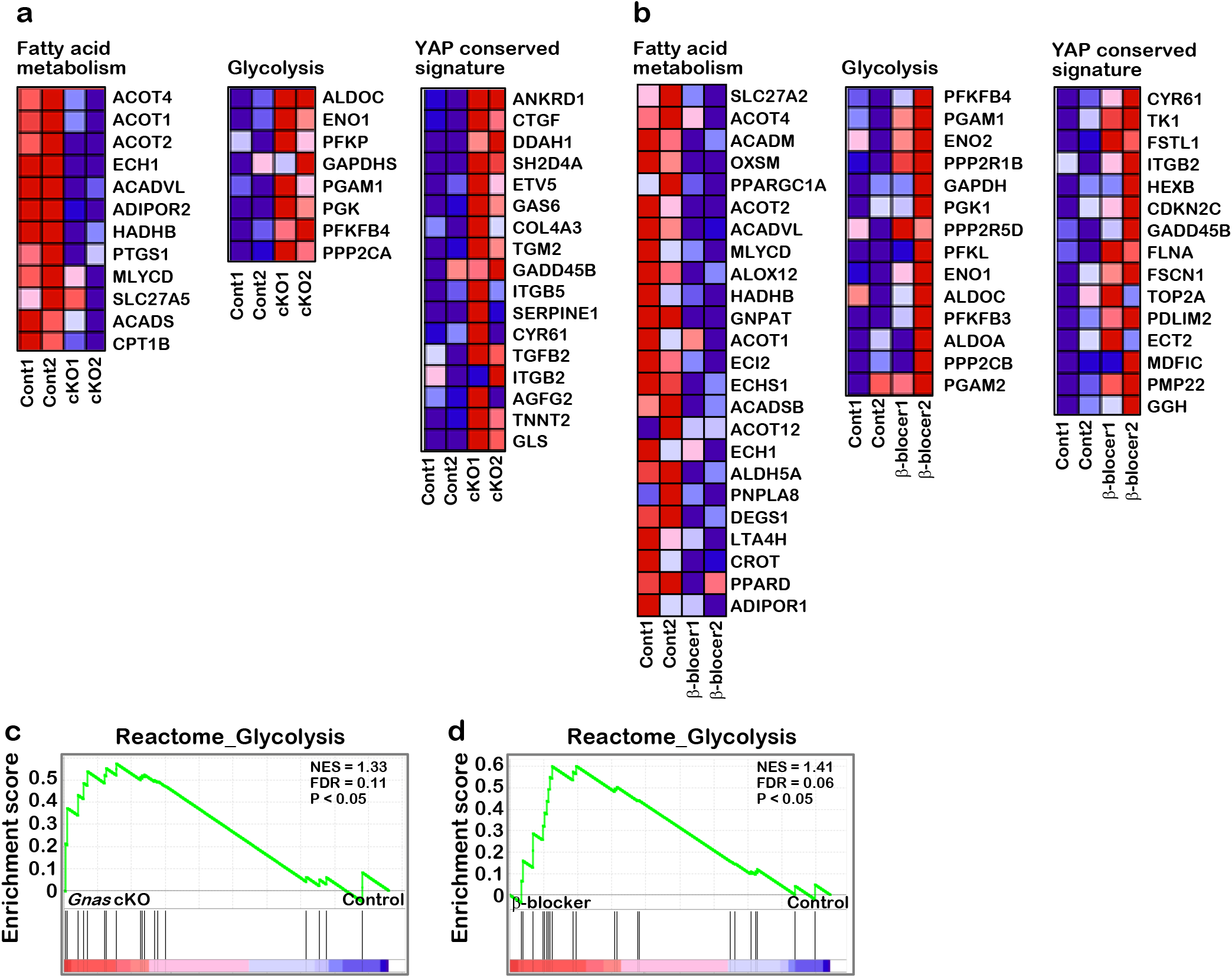
Differential gene expression in *Gnas* cKO and β-blocker treated hearts. **(a)** Heat map of fatty acid metabolism, glycolysis, and YAP signature related gene expression from RNA-seq data, Control vs. *Gnas* cKO hearts. **(b)** Heat map of fatty acid metabolism, glycolysis, and YAP signature related gene expression from RNA-seq data, Control vs. β-blocker treated hearts. **(c)** GSEA identified significant enrichment of glycolysis related gene expression in *Gnas* cKO hearts. **(d)** GSEA identified significant enrichment of glycolysis related gene expression in β-blocker treated hearts.

**Supplemental Figure 5.**
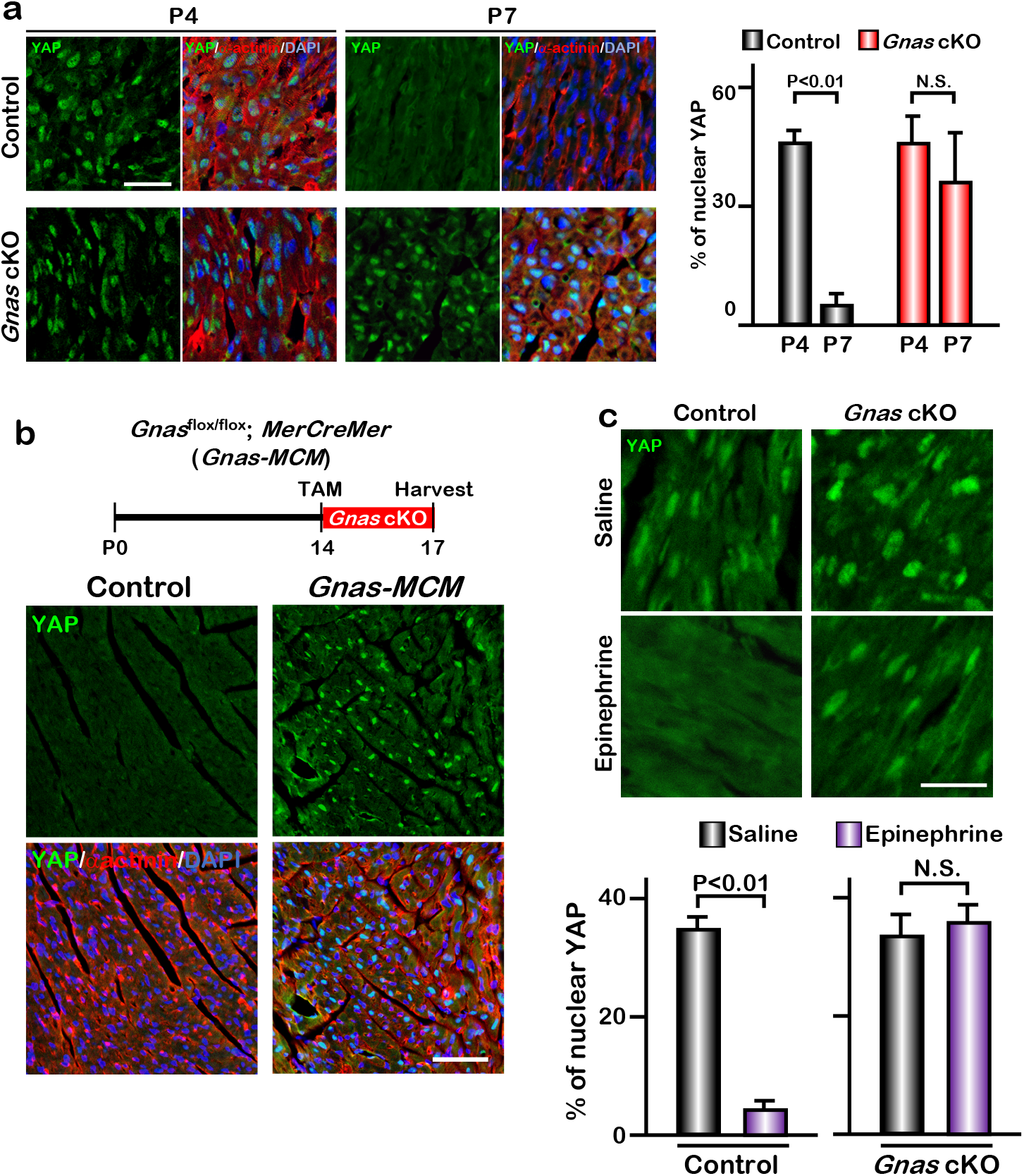
YAP activity is regulated by Gαs. **(a)** YAP and cardiac α-actinin immunostaining of P4 and P7 control and *Gnas* cKO heart sections (left panel). Scale bar: 25μm. Quantification of nuclear YAP localization in cardiomyocytes of control and *Gnas* cKO hearts (right panel). Data are as mean ± SD, n=3. **(b)** Schematic illustration of experimental design. Tamoxifen was injected into *Gnas-MCM* mice at P14, and hearts were harvested at P17 (upper panel). YAP and cardiac α-actinin co-immunostaining of heart sections from control and *Gnas* cKO hearts at P17 (lower panel). Scale bar: 50μm. **(c)** YAP immunostaining of P7 WT and *Gnas* cKO hearts treated with saline or epinephrine. Scale bar: 25μm. Quantification of nuclear YAP localization in cardiomyocytes of P7 WT and *Gnas* cKO heart treated with saline or epinephrine (lower panel). Data are as mean ± SD, n=3. N.S., not significant.

**Supplemental Figure 6.**
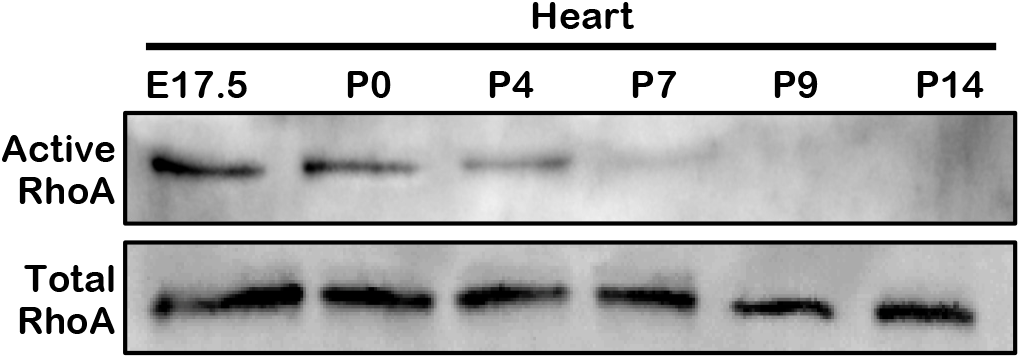
RhoA activity is only detected in the embryonic and neonatal hearts. Active-RhoA pull-down assay of wild type hearts at several time points.

**Supplemental Figure 7.**
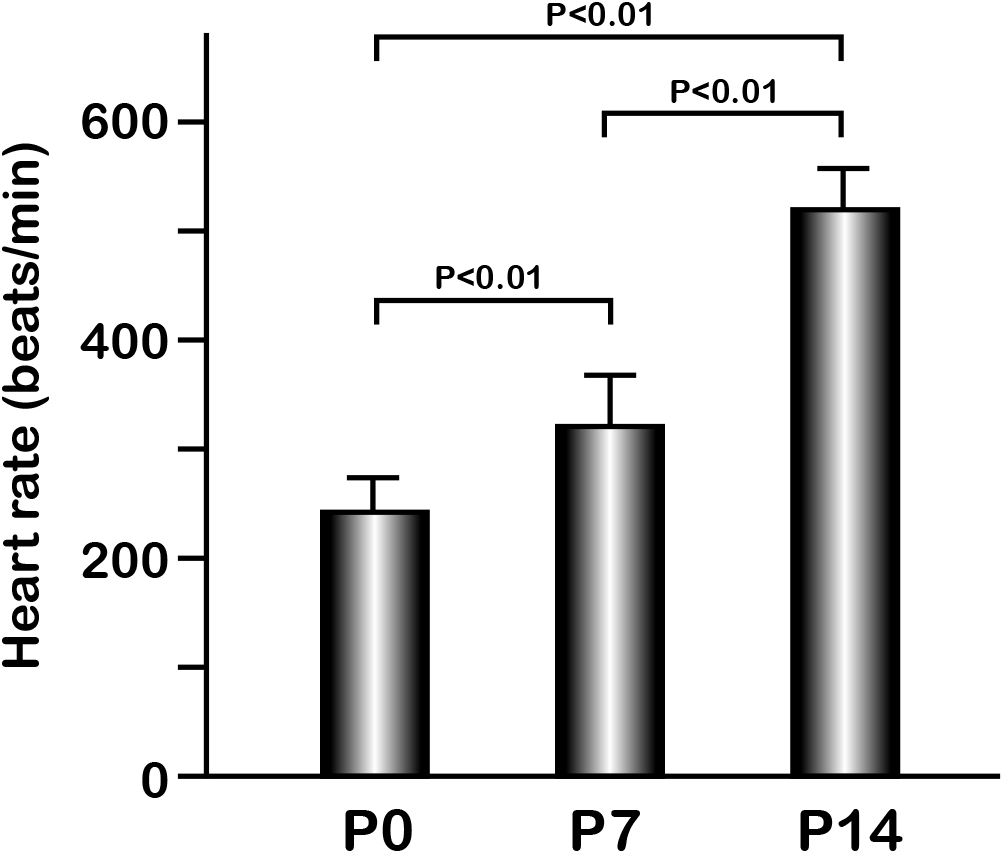
Comparison of heart rate at different ages. Heart rate at P0, P7, and P14 in C57BL6 mice. Data are as mean ± SD, n=5.

